## Supplementary information for "Global comparative structural analysis of responses to protein phosphorylation"

### Supplementary information: Global comparative structural analysis of responses to protein phosphorylation

#### 1 Supplementary figures

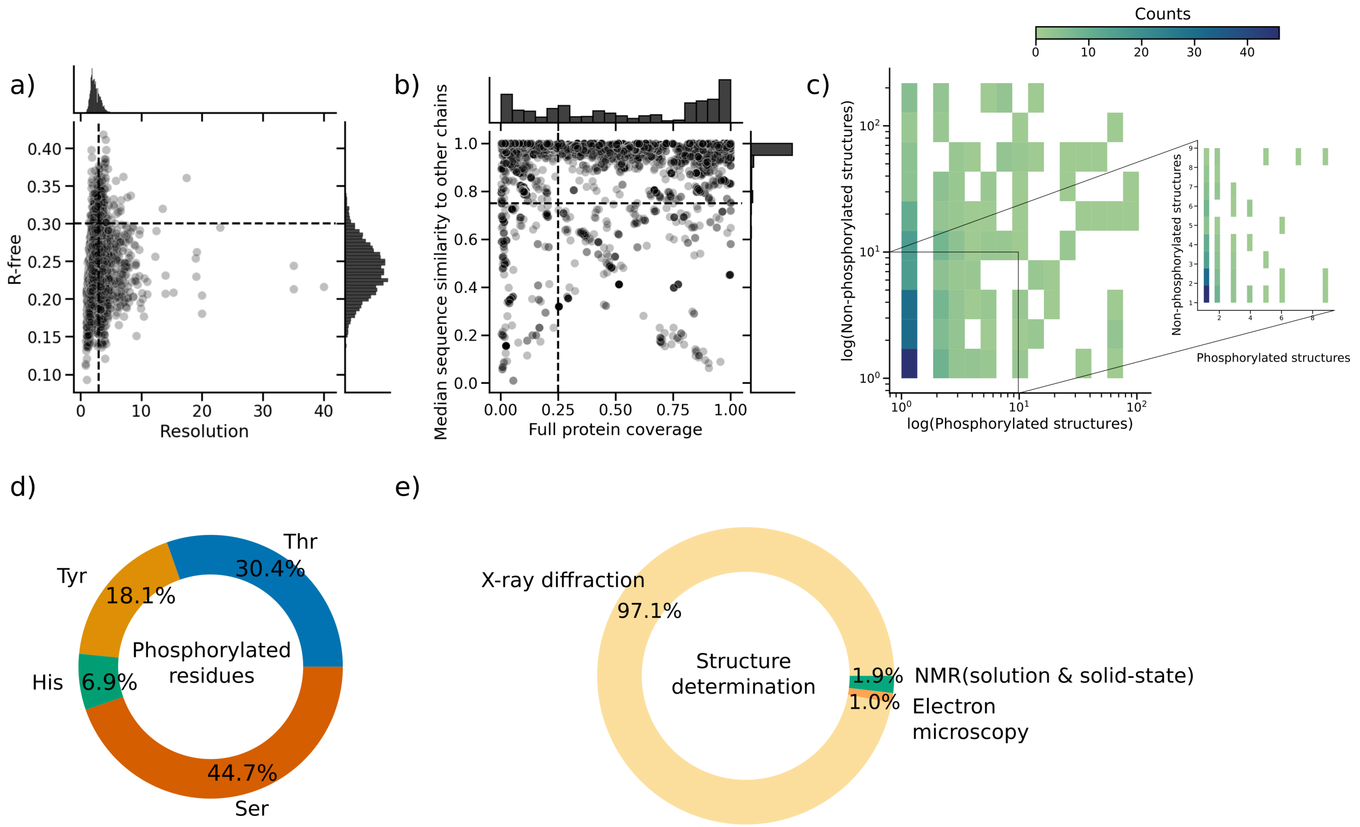


Figure S1: Curation of structural data. a) Scatterplot of structural resolution versus R-free for each retrieved structure. Dashed lines indicate selected cutoff points. b) Scatterplot of structural coverage (relative to the full UniProt canonical sequence) versus the median sequence similarity to all other structures of the same protein in the dataset. The latter serves as a metric of overlap. Dashed lines indicate selected cutoff points. c) 2D histogram showing the number of phosphorylated structures versus the number of non-phosphorylated structures for each protein in the filtered dataset. The inset focuses on the range from zero to 10 structures. d) Donut chart displaying the distribution of different phosphorylated residues in the filtered dataset. e) Donut chart illustrating the methodologies used to solve the structures in the filtered dataset.

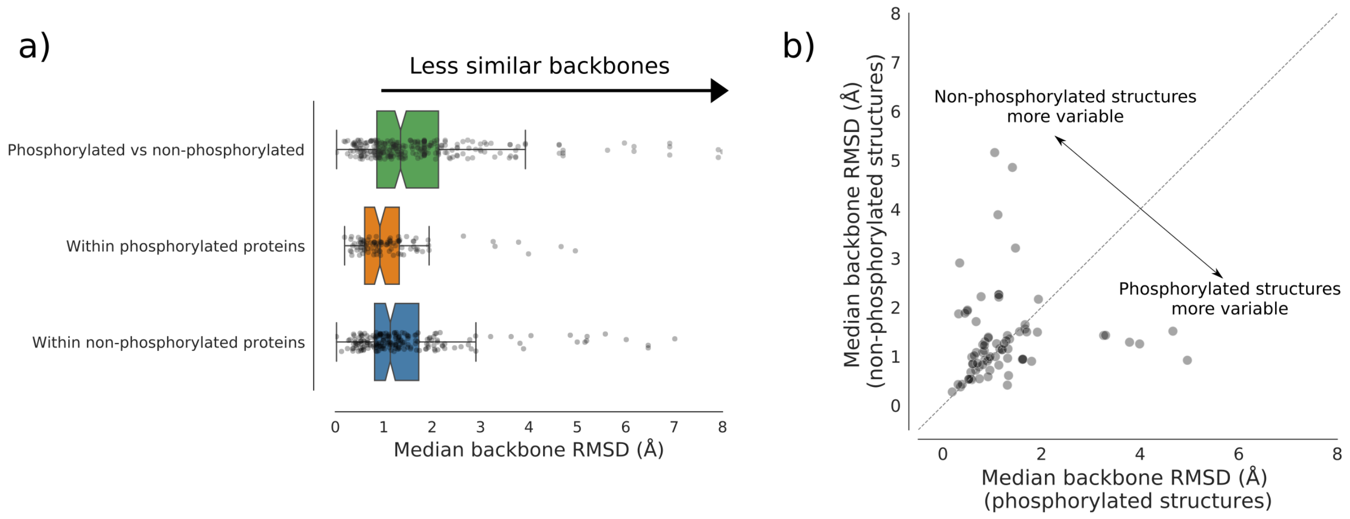


c)

| Median backbone RMSD | Excluding comparisons between structures with the same lead author: No | Yes |
|---|---|---|
| Phosphorylated vs non-phosphorylated | 1.28 ± 3.13 | 1.35 ± 2.90 |
| Within phosphorylated proteins | 0.61 ± 1.56 | 0.92 ± 1.12 |
| Within non-phosphorylated proteins | 0.90 ± 1.66 | 1.14 ± 1.81 |

Figure S2: Investigating the influence of the lead author effect on observed conformational changes. To control for possible batch effects due to structures being produced by the same research lab, we excluded all pairwise comparisons of structures with the same lead a uthor. a) Boxplot of conformational changes between phosphorylated and non-phosphorylated structures, and structural variability within each set. Points represent median changes per phosphosite. Notches indicate 95% confidence intervals. b) Scatterplot comparing median backbone RMSD per phosphosite within phosphorylated structures (x-axis) to non-phosphorylated counterparts (y-axis). The diagonal line represents the identity line; 59% of points are above the line. Only includes phosphosites with at least two structures in each state. After excluding comparisons betweeen structures sharing the same lead author, 78 such phosphosites are available. c) Table showing the median backbone RMSD for different comparisons over the whole dataset, either before excluding comparisons between structures sharing the same lead author (left) or after (right).

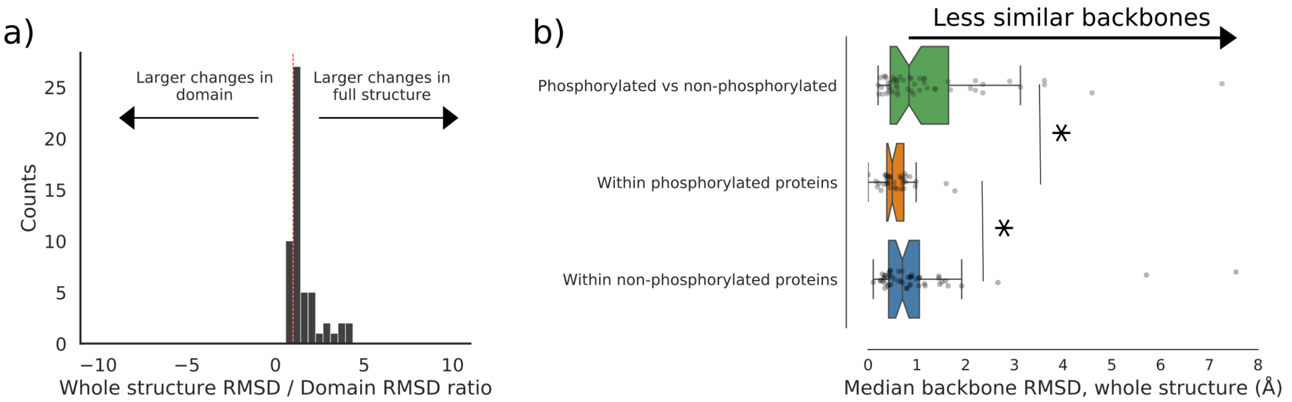


Figure S3: Comparison of the effect of phosphorylation on protein backbones in protein domains versus the whole structure. a) Histogram of the ratio of backbone RMSD in the whole structure to backbone RMSD only in the phosphorylated Pfam domain. A ratio greater than 1 indicates greater changes in the overall structure than in the phosphorylated domain. b) Boxplot of conformational changes between phosphorylated and non-phosphorylated structures (using the whole structure), and structural variability within each set. Points represent median changes per phosphosite. Notches indicate 95% confidence intervals; outliers beyond 8Ånot shown.

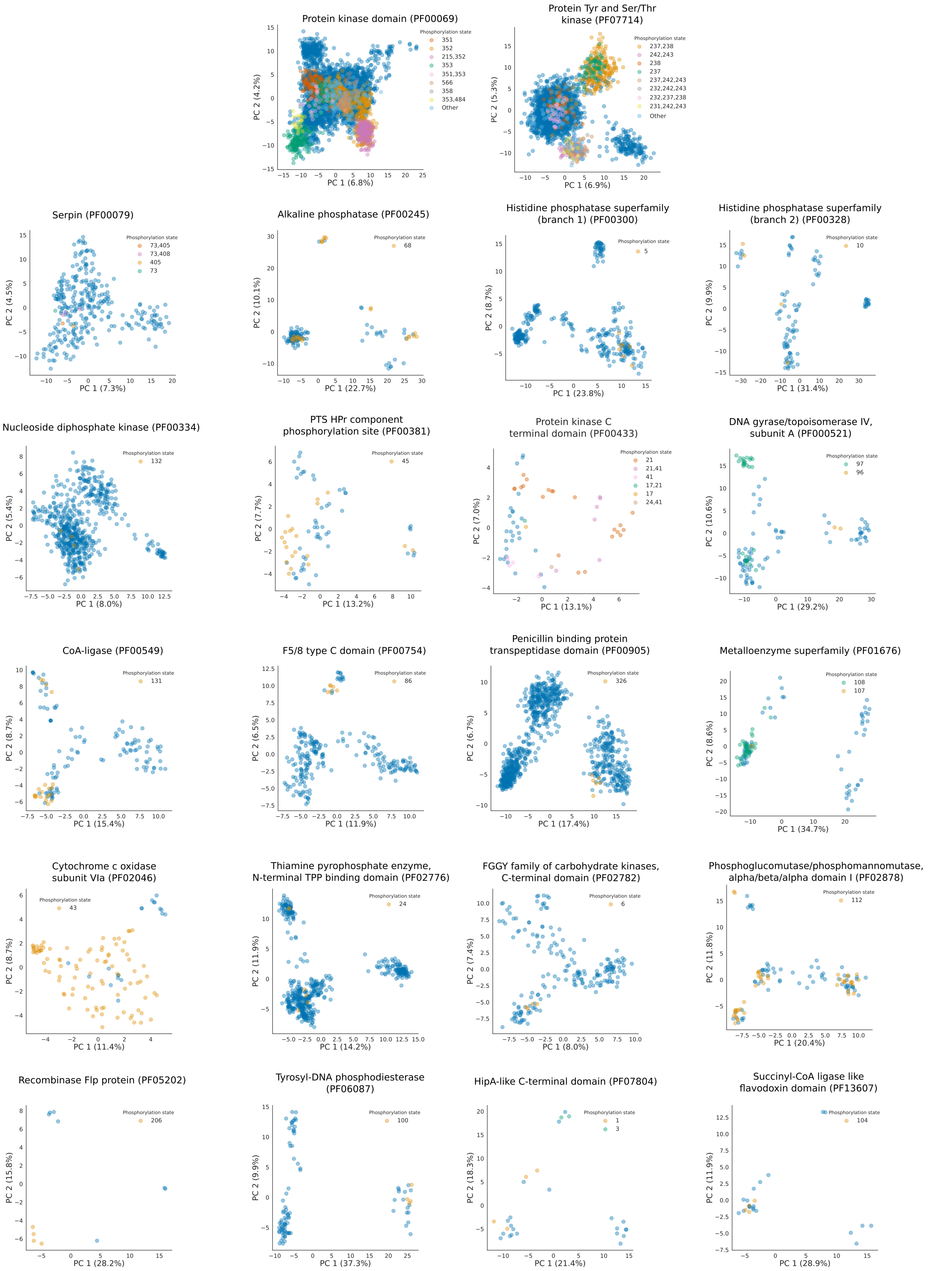


Figure S4: Additional examples of low-dimensional, domain-specific conformational landscapes. Blue dots represent non-phosphorylated structures, while other colors indicate specific phosphorylation states. For clarity, due to the high number of different phosphorylation states observed in the protein kinase domain (PF00069) and protein tyrosine and serine/threonine kinase (PF07714; both displayed on top), only the 8 most common states (besides non-phosphorylated proteins) are colored distinctly. Phosphosite indexes in the legend are assigned based on sequence alignments of the analyzed structures.

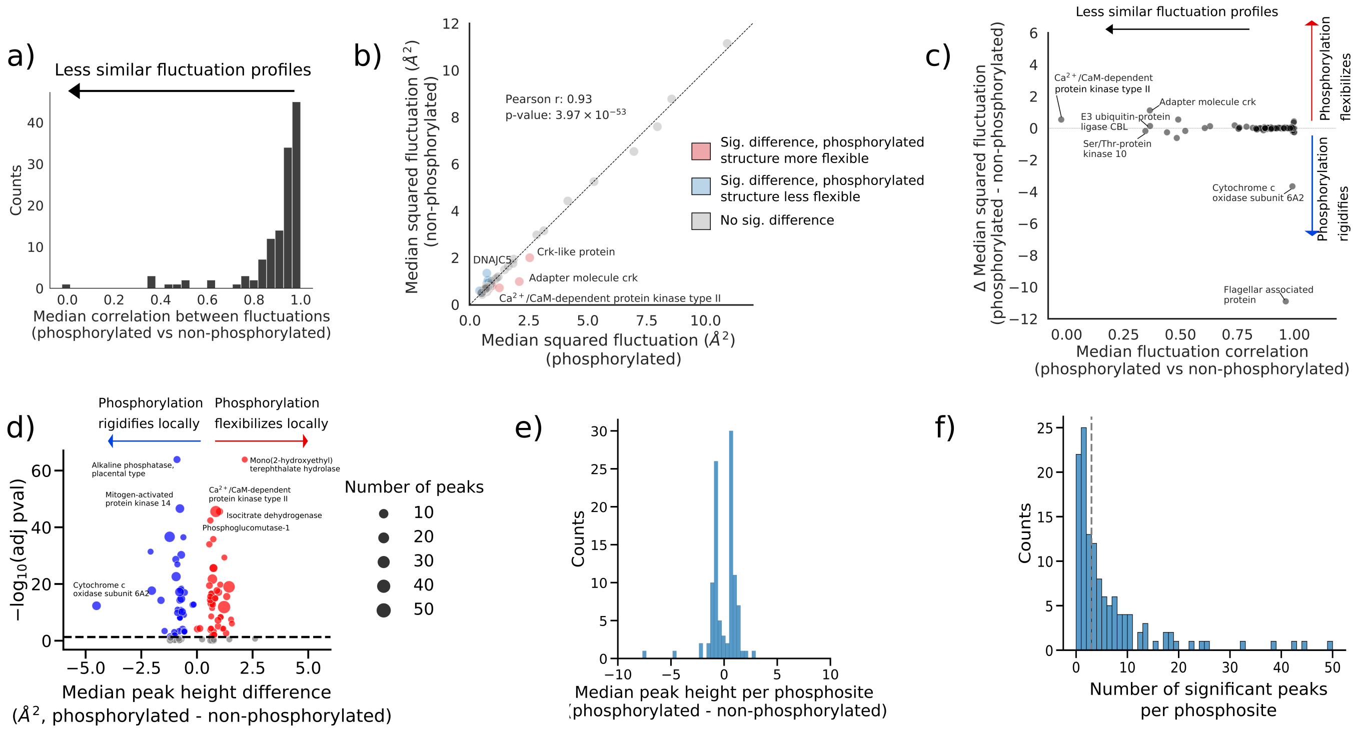


Figure S5: Changes in predicted global and local residue fluctuations. a) Histogram showing the median correlation in residue fluctuation profiles between phosphorylated and non-phosphorylated structures. b) Scatterplot comparing the median fluctuations of phosphorylated structures (x-axis) to their non-phosphorylated counterparts. Among cases with significant changes, ~47% showed increased and ~53% showed decreased flexibility. DNAJC5 refers to DnaJ homolog subfamily C member 5. Outliers beyond 12 Å are excluded. c) Scatterplot comparing the median correlation in residue fluctuations between phosphorylated and non-phosphorylated structures (x-axis) with the difference in median protein-wise fluctuation between phosphorylated and non-phosphorylated proteins (y-axis). d) Scatterplot illustrating the magnitude and significance of local changes in dynamics. Bubble size proportional to the number of significant local peaks identified. e) Histogram of the number of significant local peaks (i.e., differences in residue fluctuations exceeding a specified threshold) per phosphosite. The dashed line indicates the median number of peaks (three). f) Histogram of the median height of each peak per phosphosite. Overall negative values indicate greater fluctuations in the non-phosphorylated protein; overall positive values, the reverse.

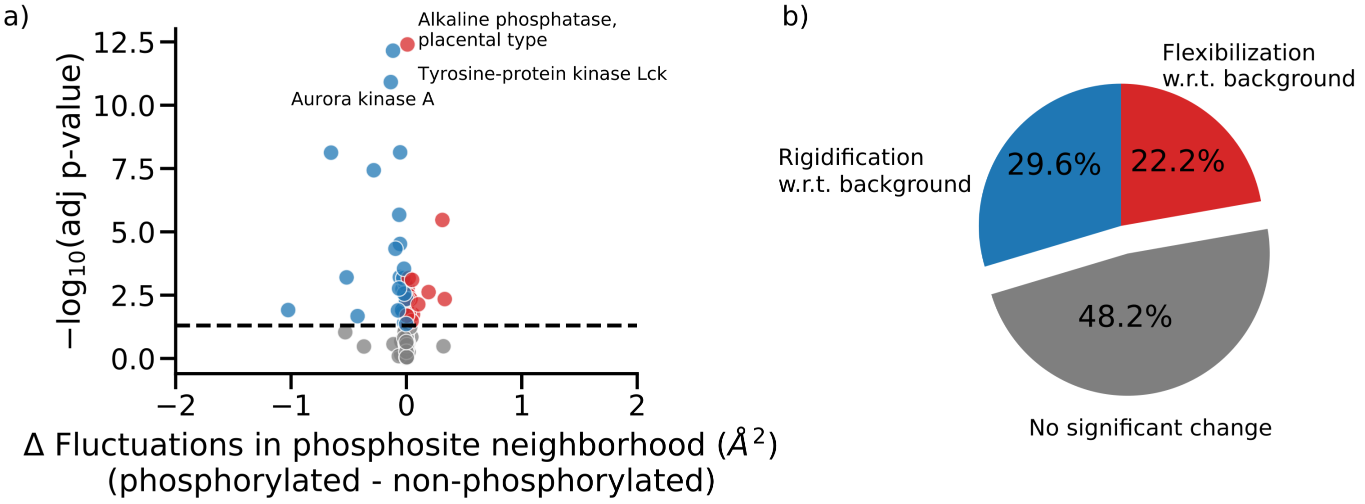


Figure S6: Local changes in predicted dynamics in the neighborhood of the phosphosite with respect to the rest of the protein. 51.8% of phosphorylation events are associated with such changes, which are typically extremely minor (median change with respect to background: $-7.6 \times 10^{-3}$ Å$^2$, among significant cases) and with no consistent direction of the effect. a) Scatterplot illustrating the magnitude and significance of changes in dynamics in the phosphosite neighborhood with respect to the background upon phosphorylation. b) Pie chart categorizing the types of local changes in dynamics in the phosphosite neighborhood.

a)

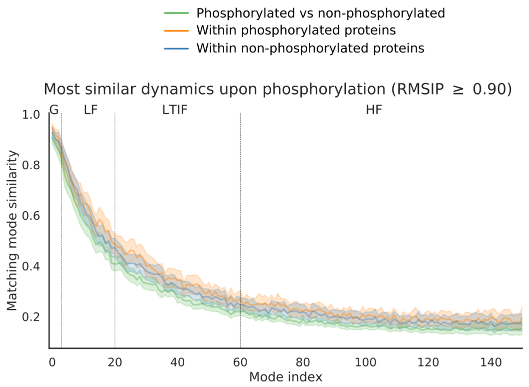


| | |
|---|---|
| N | 100 |
| Adj. R squared | 0.419 |
| F-statistic | 12560 |
| p-value | <0.001 |

| | **Coefficient** | **p-value** |
|---|---|---|
| Intercept | 0.636 | <0.001 |
| Comparison between groups | -0.0299 | <0.001 |
| Comparison within phosphorylated | 0.016 | <0.001 |
| sqrt(Mode index) | -0.0395 | <0.001 |

b)

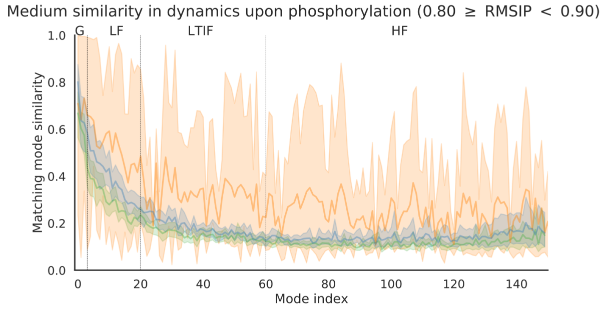


| | |
|---|---|
| N | 18 |
| Adj. R squared | 0.259 |
| F-statistic | 679.8 |
| p-value | <0.001 |

| | **Coefficient** | **p-value** |
|---|---|---|
| Intercept | 0.371 | <0.001 |
| Comparison between groups | -0.035 | <0.001 |
| Comparison within phosphorylated | 0.192 | <0.001 |
| sqrt(Mode index) | -0.022 | <0.001 |

c)

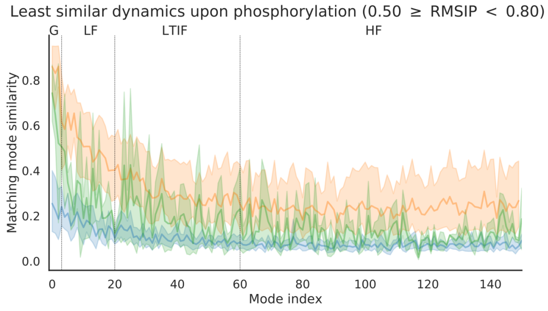


| | |
|---|---|
| N | 10 |
| Adj. R squared | 0.195 |
| F-statistic | 358.8 |
| p-value | <0.001 |

| | **Coefficient** | **p-value** |
|---|---|---|
| Intercept | 0.252 | <0.001 |
| Comparison between groups | -0.053 | <0.001 |
| Comparison within phosphorylated | 0.111 | <0.001 |
| sqrt(Mode index) | -0.008 | <0.001 |

Figure S7: Comparison of normal modes in different bins of dynamics similarity, from most similar (top) to least similar (bottom). Left column, similarity of matching pairs of modes over different frequency regimes. Bands indicate 95% confidence intervals. Right column, summary of linear regressions modelling the relationship between mode similarity for each bin. All regressions were statistically significant, as were the model coefficients, indicating significant differences in normal modes regardless of the bin.

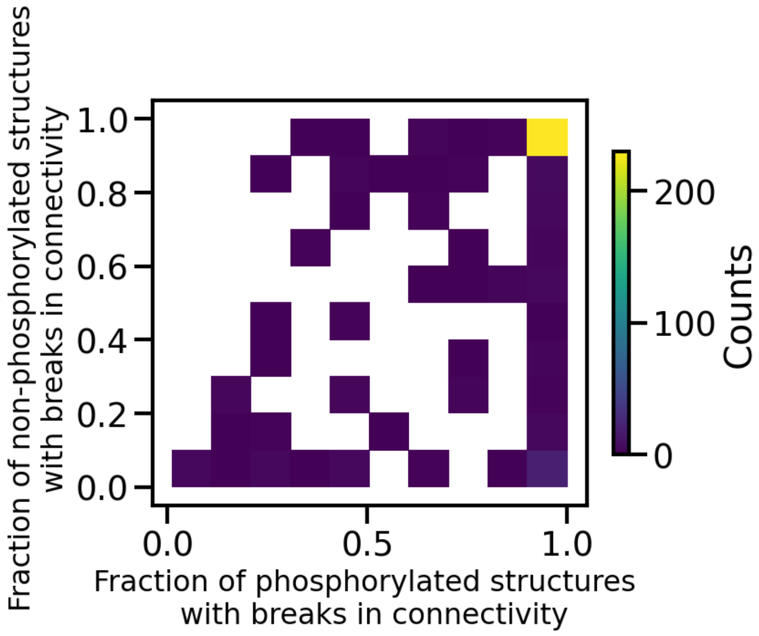


Figure S8: 2D histogram showing the number of phosphorylated and non-phosphorylated structures that were filtered out (for all 347 unique phosphosites) for normal mode analysis due to breaks in connectivity.

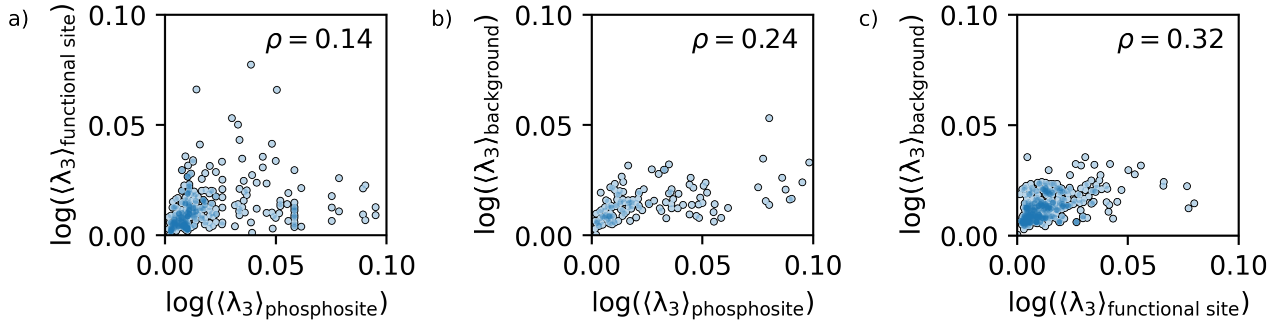


Figure S9: Correlation analysis of the mean extensive stretches between three groups: phosphosites (± 2 residues), functional sites, and background residues. Although the observed correlations were consistently relatively low, they were nonetheless statistically significant compared to the null hypothesis of uncorrelated, normally distributed data ($p = 0.001$, $p = 0.003$, and $p = 6 \times 10^{-12}$, respectively).

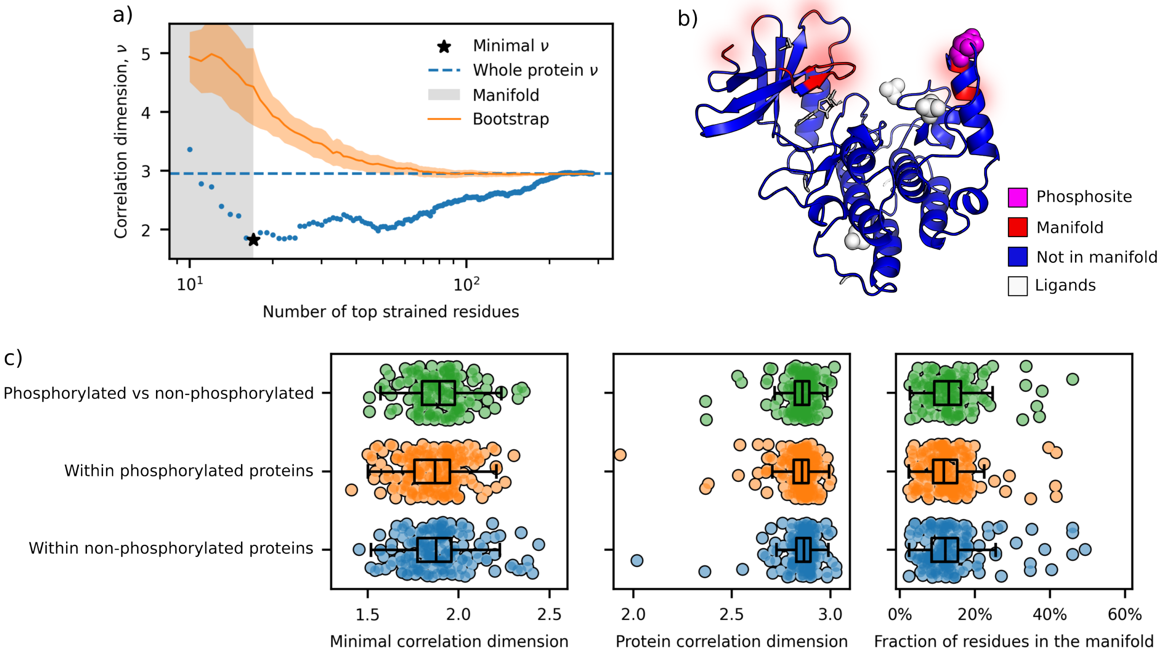


Figure S10: Strain manifold analysis reveals a conserved structural architecture between phosphorylated and non-phosphorylated proteins. a) Example of manifold determination using the correlation dimension $\nu$ for residues ranked by strain (PDBs 7P7F_B vs. 4HNF_A). The correlation dimension values are plotted as a function of the number of top-strained residues, with the minimum (denoted by a star) indicating the residues belonging to the high-strain manifold. Bootstrapping ($n = 40$) demonstrates that randomly distributed strained residues do not form a low-dimensional manifold. b) Spatial visualization of residues within the strain manifold, where strain is concentrated in the upper region of the structure. c) General properties of high-strain manifolds remain consistent across comparisons between different groups, with no significant differences ($p > 0.05$, Kruskal-Wallis test) in manifold dimension, overall protein correlation dimension, or the proportion of residues within the manifold.

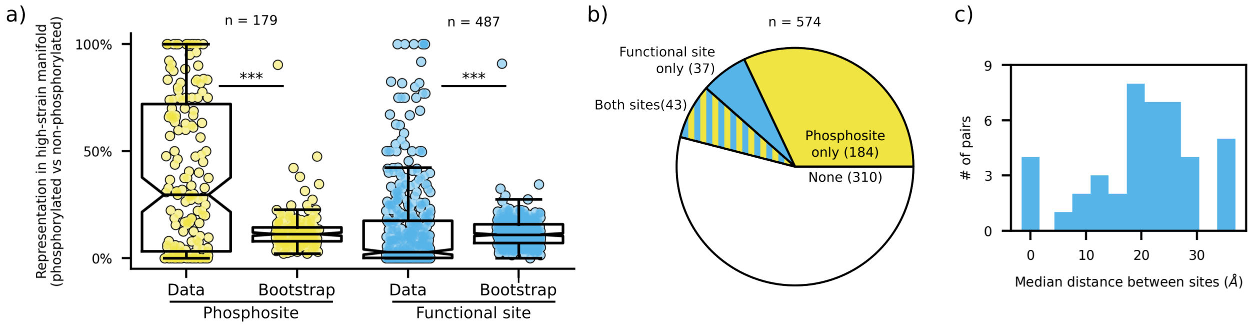


Figure S11: Representation of phosphosites and functional sites in the high-strain manifold. a) Boxplot of the representation of phosphosites and functional sites in the manifold, averaged across all pairwise comparisons per phosphosite. Bootstraps were generated by randomising the strain distribution over different residues ($n = 50$). Statistical significance was determined using two-tailed Wilcoxon signed-rank tests (*<0.05, **<0.01, ***<0.001). b) Share of pairs of phosphosite / functional sites with high representation in the manifold. We identify each site as significantly represented by comparing them individually to the bootstraps of the same protein when satisfying the following criterion: $\langle R \rangle^{\text{data}} > \frac{1}{n}\sum_i^n \langle R \rangle_i^{\text{bootstrap}} + 3\sigma^{\text{bootstrap}}$, where $\sigma$ is the standard deviation of the representation in the bootstrap samples. c) Histogram of distances between phosphosite and functional site pairs identified to belong to the high-strain manifold.
